# Improved EPR sensitivity for aqueous biological samples using low-volume multi-channel cells and dielectric resonators

**DOI:** 10.1101/2025.06.03.657726

**Authors:** Richard R. Mett, Alexander M. Garces, Anand Anilkumar, Joseph T. Wehrley, Michael T. Lerch, Candice S. Klug, Jason W. Sidabras

## Abstract

Reducing sample volumes for electron paramagnetic resonance (EPR) spectroscopy applications places increasing demands on hardware design to preserve or enhance EPR signal intensity. In this work, the design, fabrication, and testing of dielectric resonators and multi-channel aqueous sample cells for applications in X-band (nominally 9.5 GHz) EPR is presented. Our aim was to maximize the EPR signal intensity for sample sizes of 3–4 μL and 200 nL. These advances are summarized as follows: single-crystal sapphire and rutile dielectric resonators with very low loss tangent and high resonator efficiency; minimum dielectric resonator coupling to radiation shield to reduce ohmic losses; 3D printed aqueous sample cells with thin multi-channel construction to minimize radio-frequency dissipation in the sample; and a Gordon coupler for maximum coupling range and minimum stored energy to eliminate frequency shifts during tuning. Sample cell geometries were designed by leveraging insights gained from analytic theory to inform finite-element modeling of electromagnetic fields. Experimental comparisons of multi-channel sample cells using a sapphire resonator exhibited a 2.2-fold increase in EPR signal intensity compared with a standard capillary at 3–4 μL, while simulations predict an additional 23% improvement with further 3D printing advances. For samples at 200 nL, a rutile dielectric resonator with a multi-channel sample cell was simulated to improve EPR sensitivity by a 2.7-fold increase compared with a capillary at the same volume.

## I. INTRODUCTION

Electron paramagnetic resonance (EPR) spectroscopy is a critically important technique in biomedical research with a unique ability to detect naturally occurring or engineered unpaired electrons in complex biological environments. EPR has wide-ranging applicability to structural biology, metalloprotein research, redox biology, rational drug design, and clinical diagnostics [1–13]. Sample volume and concentration requirements are a critical consideration in all biomedical EPR studies. For example, many biomedical EPR samples require low concentrations to avoid aggregation or other types of instability; difficult-to-express recombinant proteins and patient- or animal-derived samples are often not obtained in sufficient amounts to study by EPR, and high-volume kinetic measurements or large drug screens by EPR are sample limited. Here, we aim to lower the required spin concentrations for data acquisition while maintaining low sample volume requirements to enable the broader use of powerful continuous-wave (CW) EPR spectroscopy technologies in the biomedical sciences.

Froncisz and Hyde made the novel leap to low-volume samples (i.e., approx. 2 µL) in 1982 with the development of the loop-gap resonator (LGR) [14]. In 2019, we made the next state-of-the-art step by developing a dielectric LGR (dLGR) to enable even smaller volumes (2–0.5 µL) without compromising EPR signal intensity [15]. Here, we capitalize on these advancements with a dielectric resonator (DR) coupled with innovative sample tube geometries to further reduce the number of spins required per sample while maintaining similar EPR sensitivity.

To understand the relationship between sample volume and resonator characteristics, one can show that the EPR signal voltage derived in Feher’s fundamental paper on EPR sensitivity considerations [16], for spins that are saturated, is proportional to the product of the resonator efficiency,

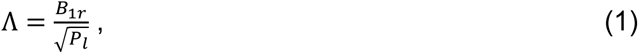

and the sample volume [17]. In Eq. (1), *B*_1*r*_ represents the maximum available radio frequency (RF) magnetic field for a given incident microwave power and *P*_*l*_ represents the power dissipation in the resonator. Consequently, when the sample volume is fixed, the EPR signal is maximized by maximizing Λ. Primarily, there are two ways to increase the resonator efficiency: decreasing resonator size and decreasing the total RF losses from both the sample and resonator. This is captured quantitatively by the following expression, which can be derived from the stored energy definition of the quality factor *Q*-value of a resonator,

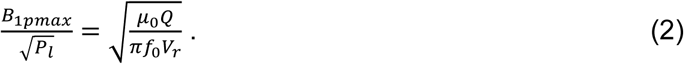

In this equation, we have the maximum peak magnetic field value in the resonator *B*_1*pmax*_,^1^ the unloaded (eigenmode) *Q*-value, the resonance frequency *f*_0_, the vacuum permeability *μ*_0_, and the effective resonator volume defined by

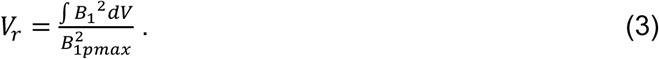

In Eq. (3), the integral is over the entire resonator volume.^2^ Equation (2) shows that at fixed frequency, maximum Λ occurs at maximum *Q*-value and minimum resonator volume. At X-band, cavity resonators have a relatively low Λ-value of about 0.1 mT/W^1/2^ because even though the *Q*-values are high, in the several thousands, *V*_*r*_ is large. LGRs have 3–6 times the Λ-value of cavities despite having much lower *Q*-values, in the hundreds, because LGRs have much smaller volume. In the present study, we use single-crystal (SC) dielectrics of very low loss that significantly reduce the size of the DR to achieve both high Λ-values with high *Q*-value. We focus on two sample volume considerations: *(i)* low microliter volumes (3–4 μL) for conventional EPR and flow experiments, and *(ii)* nanoliter volumes (200 nL) for microfluidics and high-throughput EPR. Ultra-high purity synthetic SC sapphire was used for the DR for the 3-4 µL sample size, and ultra-high purity synthetic SC rutile DR was used for the 200 nL sample. The volumes of the resulting DRs permit sample insertion without significant deleterious effects. Our designs leverage insights gained from three previous DR studies from our laboratory [15, 19, 20] as described in Sec. III below.

Insertion of an aqueous sample into a DR can decrease the Λ-value for three reasons: *(i)* RF power losses in the aqueous sample can be significant, lowering the resonator *Q*-value; *(ii)* the sample cell material can have RF losses, and *(iii)* the sample access hole in the DR increases the effective resonator volume (at fixed frequency) when the sample and sample cell material together have a lower effective dielectric constant than the DR. Of the three causes above, sample losses can be the most significant, particularly if a simple capillary tube is used. Sample tube geometries beyond a cylindrical tube shape were introduced previously by our group in the form of the Θ-geometry (cylindrical tubing with a septum) for a 1 mm sample diameter LGR applications and Aquastar designs for a five-loop–four-gap LGR and cylindrical cavity resonators [21, 22]. The first-generation sample cell geometries required extrusion techniques and thus limited them to larger sample volumes and prevented further advances in, and implementation of, low volume sample tube geometries. Advances in 3D printing technologies and resin development now allow for an opportunity to continue the pursuit of complex sample cell designs on the micrometer scale for the biomedical EPR community.

To accommodate the biomedical needs of increased EPR signal intensity and to provide the broader scientific community with accessible technologies, we capitalized on the development of the DR concept by optimizing these resonator technologies and sample cell geometries to increase sensitivity beyond that achieved upon introduction of the LGR nearly four decades ago [14]. Based on previous work from our laboratory [21, 23–26], here we report designs of thin multi-channel sample cells that have much lower RF losses for the same sample volume than a capillary tube. New methods of lowering RF losses in both the water and sample cell are also reported, and we discuss the design and fabrication of the Gordon coupler, which is also based on previous work from this laboratory [27]. Fabrication and EPR characterization of the sapphire resonator are presented, showcasing the experimental improvement gained through these advances.

## II. ANALYSIS TOOLS

Analytic calculations were performed using Wolfram Mathematica (Champaign, IL) ver. 12.3.1. Finite-element simulations were performed using Ansys Electronics Desktop 2021/R2 High Frequency Structure Simulator (HFSS) (Canonsburg, PA) running on an HP Elitebook 850 G8 laptop with Intel Core i7 1165G7 / 2.8 GHz (4 cores, 4.7 GHz boost) 16 GB RAM and 512 GB SSD. Eigenmode solutions were used to simulate the designs providing the resonance frequency and *Q*-value without the requirement to couple the system. Designs of the DRs with the radiation shield and designs of the multichannel sample cells were all done using eigenmode solutions. Critical coupling to the waveguide was also simulated using eigenmode solutions. The iris size and Gordon coupler position are adjusted such that the *Q*-value of the coupled resonator is half the value of the *Q*-value of the resonator with no iris (i.e., the iris is filled with conductor). Final designs were fully simulated in driven mode with a solution frequency equal to the eigenmode frequency, and a narrow frequency sweep was used to verify critical coupling and determine the coupling (β).

For each resonator configuration, the HFSS fields calculator was used to determine the peak Λ-value and the saturable EPR signal strength (*S*_*s*_) using the methods and equations described in Mett et al. [15]. The equations for Λ and *S*_*s*_ are

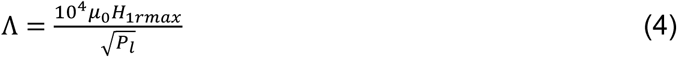

and

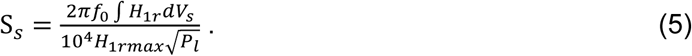

In these equations, *H*_1*r*_ is the rotating RF magnetic field component in the sample and max indicates the maximum value of the sample volume. The integral is done over the sample volume. The power losses include the metallic walls, dielectric, sample, and sample cell. Equations (4) and (5) were developed in [23, 24, 28].

The birefringence of the SC sapphire and SC rutile was simulated in ANSYS HFSS by setting the appropriate values of the dielectric tensor (relative dielectric constant and loss tangent) for each of the three spatial dimensions of the crystal. The relative dielectric constant of sapphire is 11.59 parallel to the c-axis (the (0 0 1) plane) and 9.40 perpendicular, and the loss tangent is 6.5 × 10^−6^ parallel to the c-axis and 1.8 × 10^−5^ perpendicular [29]. The relative dielectric constant of SC rutile is 165 parallel to the c-axis and 86 perpendicular, and the loss tangent is 10^−4^ parallel to the c-axis and 8.5 × 10^−5^ perpendicular [30].

Other materials were analyzed as homogeneous: a water sample at 20 °C has a relative dielectric constant of 61.61 and loss tangent 0.5184 [31], PTFE (polytetrafluoroethylene) has a relative dielectric constant of 2.08 and a loss tangent of 3.7 × 10^−4^ [32], Rexolite^®^ has a relative dielectric constant of 2.54 and a loss tangent of 4.7 × 10^−4^ [32], HTL photopolymer (Boston Micro Fabrication, Maynard, MA) has a relative dielectric constant of 3.45 and a loss tangent of 0.0245 [33].

## III. DIELECTRIC RESONATOR AND RADIATION SHIELD

Shown in Fig. 1 is the designed and fabricated geometries for the DR and radiation shield, along with the Gordon coupler. The designs of the DRs and shield are based on results of Mett et al. [15, 19, 20]. In Ref. [19], it was found that the interaction between a DR of high relative dielectric constant and a surrounding conducting cylindrical TE_011_ cavity resulted in a mode splitting in which the maximum Λ-value for either mode was the same as the isolated DR. In Ref. [15], it was found that for a rutile DR placed in the center of an LGR, a similar mode splitting occurred, but the maximum Λ-value resulted for only one of the two modes, namely the parallel mode, in which the RF magnetic fields of each resonator add constructively. Even in this parallel mode, the maximum Λ-value was not larger than the maximum of either resonator individually. The conclusion of Ref. [19], for high *Q*-value, states that the maximum Λ-value could only occur when there is very low inductive coupling between the DR and LGR (Fig. 4 of Ref. [15]). In this limit, the LGR inner loop diameter is significantly larger than the DR, and the LGR acts simply as a coupler to the DR.

**Figure 1:**
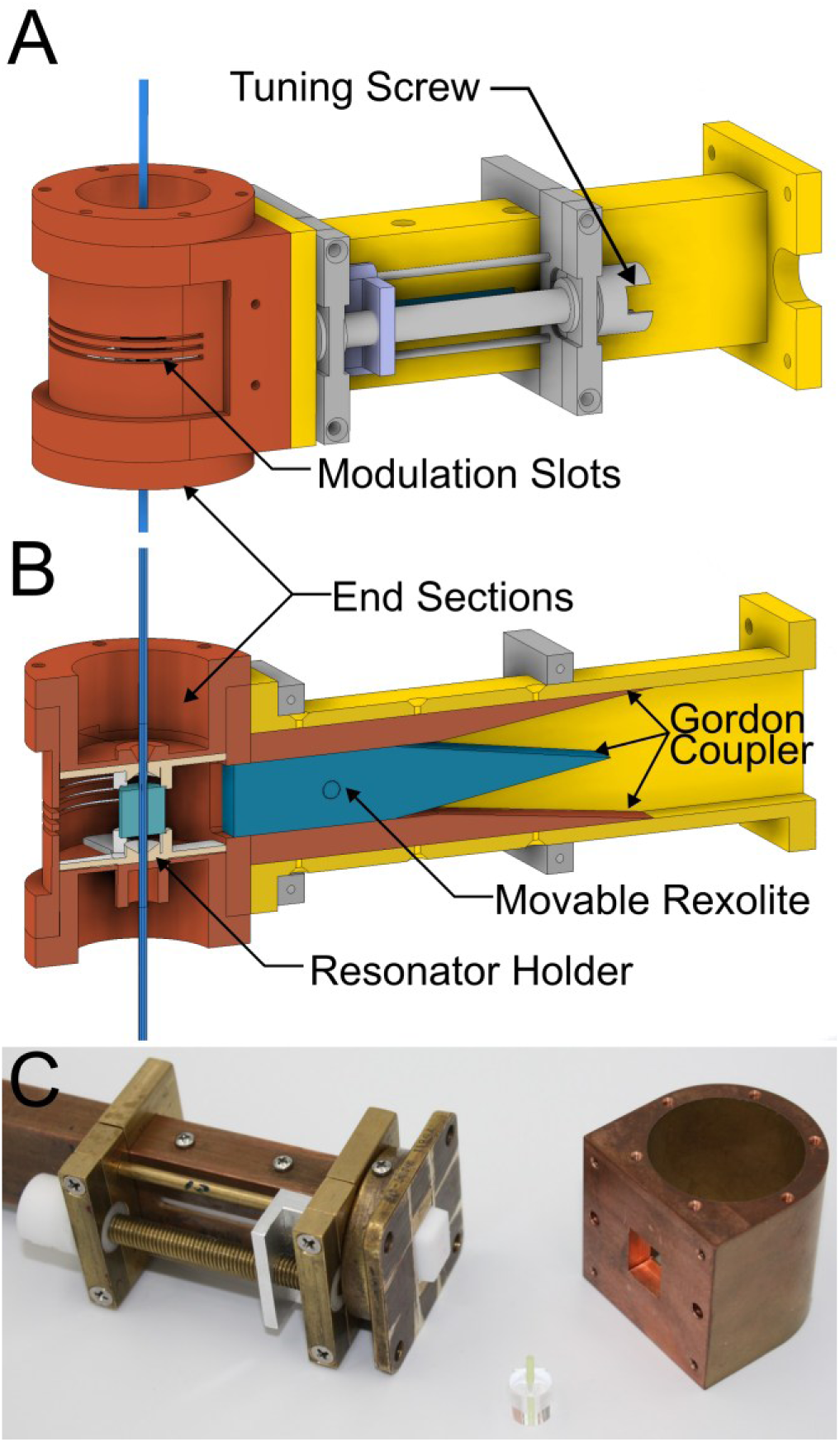
The resonant probe geometry. (A) Illustration of the assembled system highlighting the modulation slots and the tuning screw, which move the Gordon coupler for match adjustment. (B) A half-slice illustration depicting the Gordon coupler geometry and resonator holder. (C) Photograph of the disassembled resonator and shield.

From these results, maximum Λ would result under conditions of minimum interaction between the radiation shield and the DR. The design approach was as follows: The DR and shield were designed first for sapphire because the larger size of the sapphire DR compared with rutile produces a larger RF field interaction between the DR and the shield. An isolated sapphire DR was sized with equal diameter and length to be resonant with the TE_01δ_ mode at 9.5 GHz using the analytic theory of Sec. IIB of [19]. The c-axis of sapphire was chosen to be perpendicular to the DR cylinder axis to maximize Λ [15]. This orientation makes the electromagnetic mode slightly azimuthally asymmetric, and the geometrical average of the parallel and perpendicular relative dielectric constants and loss tangents can be used to make a reasonable analytic prediction. Results are shown in Table I. A cylindrical conducting (silver) TE_011_ cavity resonant at 9.5 GHz with equal diameter and length was designed and the DR placed in the center. A mode splitting was observed with a lower frequency mode in which the DR RF magnetic field was in-phase with (parallel to) the TE_011_ cavity RF magnetic field and a higher frequency mode where the two RF magnetic fields were out of phase (antiparallel). The modes were about 8.86 GHz and 10.97 GHz, with unloaded *Q*-values of 44,000 and 41,000, respectively. These *Q*-values were between that of the empty TE_011_ cavity (31,600) and that of the sapphire DR (92,450), as expected based on theory presented in Ref. [19]. To maintain the correct mode at 9.5 GHz, the following geometries are possible: *(i)* reduce the DR size with parallel magnetic field mode; *(ii)* decrease the size of the cavity acting as a shield with parallel mode; *(iii)* increase the DR size to reduce the antiparallel magnetic field mode frequency; and *(iv)* increase the cavity size to reduce the antiparallel mode. In exploring each of these possibilities, it was found that only a smaller cavity size with parallel magnetic field mode produced a larger *Q*-value with a larger Λ-value. This method also produced a lower number of nearby unwanted modes compared with the other three methods. A systematic study varying the cavity size and the DR size while maintaining a resonant frequency of 9.50 GHz showed that a 31 mm diameter and length cavity with a slightly smaller DR produced a maximum unloaded *Q*-value of 60,000 and corresponding Λ of 2.23 mT/W^1/2^. Results of several simulations are shown in Table I. Larger and smaller cavity (shield) sizes than 31 mm produced a smaller *Q*-value because the magnetic fields near the metallic walls became larger. Also shown in the table are simulations using a perfect electric conductor (PEC) for the shield to determine the effect of the conductivity. The conductivity of the shield lowers Λ by 23% because of the extremely low loss of the sapphire. A sapphire DR with the c-axis parallel to the cylinder axis was also designed for this shield (Table I).

**Table I:**
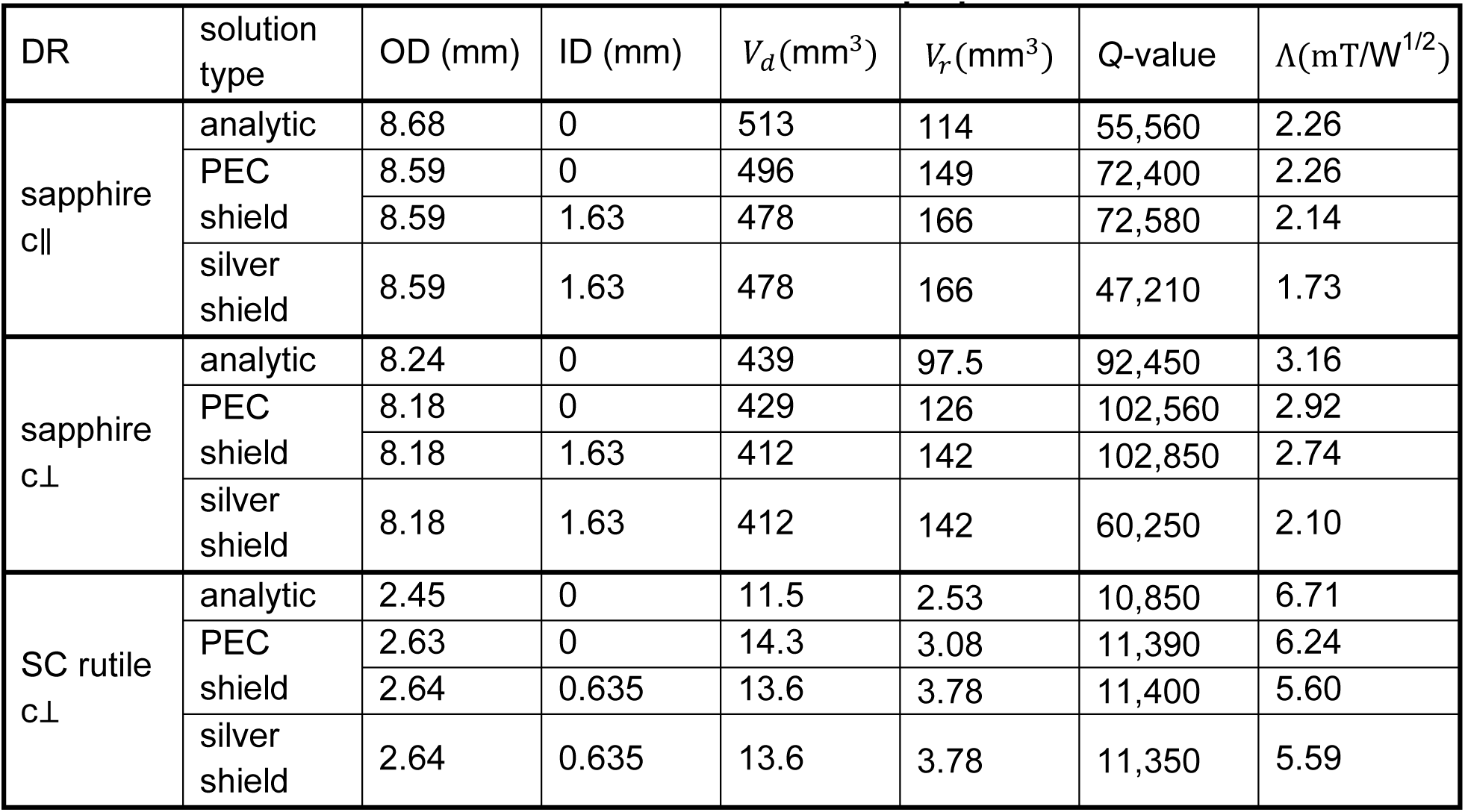
Characterization of DR properties.

In addition, an SC rutile DR with the c-axis perpendicular to the cylinder axis was sized to resonate in the same shield at 9.5 GHz and the results are shown in Table I. The conductivity of the shield has an insignificant effect on the performance of the SC rutile DR.

The outer diameter (OD) and length of the SC sapphire and rutile DRs were adjusted to be within ±6 MHz of 9.5 GHz after determining the inner diameter (ID) of the DR needed to accommodate the sample cells described in Sec. VI below. The resulting parameters are also shown in Table I. For SC rutile, there is a 10% drop in Λ due to the small size of the sample access hole.

Table I also reports the dielectric volumes *V*_*d*_ and the effective resonator volumes *V*_*r*_, Eq. (3). It is seen that, with no sample hole, the *V*_*r*_ are about 30% of *V*_*d*_ for sapphire and 22% for the SC rutile. In every case, the hole in the dielectric causes *V*_*d*_ to decrease by almost exactly the hole volume and *V*_*r*_ to increase by approximately the same amount compared with no hole. The empty hole decreases the RF magnetic field concentration and increases the effective resonator volume. In consequence, the sample hole decreases Λ, Eq. (2). The effect is more significant in rutile because the hole volume is a larger fraction of *V*_*r*_. Note also that the sample volume (which is quantitatively incorporated in Sec. VI) is about 2% of *V*_*r*_ for the sapphire and 5% for the rutile.

## IV. GORDON COUPLER

Gordon couplers were invented by J. P. Gordon in 1961 [34] and have since been implemented in numerous EPR and nuclear magnetic resonance resonators, including Mett and Hyde [27] (see also the references included therein). In Ref. [27], Gordon couplers of various materials were designed to couple a standard rectangular waveguide to an LGR with a DR inside the inner LGR loop. The Gordon coupler consists of four elements between the resonator and the standard waveguide (Fig. 1): an iris, a reduced-size waveguide section, a tapered waveguide section between the reduced-size section and the standard waveguide, and a moveable dielectric wedge that fills the reduced-size waveguide and is tapered down to zero at the standard waveguide. Gordon coupler theory is described in Sec. 2 of Ref. [27]. The essence of its operation is that the presence of the dielectric in the reduced-size waveguide permits propagation of the waveguide mode while its absence does not. The mode is evanescent in the empty part of the reduced-size waveguide: the electromagnetic fields decay exponentially with a decay length *γ*^−1^, which depends on the reduced-size waveguide dimension. Values of this and other important quantities for the rectangular TE_10_ mode and different dielectrics at 9.5 GHz are shown in Table 1 of Ref. [27]. The coupling strength is maximum when the dielectric is closest to the resonator and decreases exponentially approximately as *e*^*γz*^ with distance *z* from the resonator. Depending on the length of the reduced-size waveguide and the value of *γ*^−1^, a very large range of resonator *Q*-values can be critically coupled over a reasonable and smooth distance of travel, with larger *Q*-values monotonically matched at larger distances. Another important characteristic is that there is very low stored energy in the coupler and so the frequency pulling of the fundamental mode of the resonator is very small [27, 35]. This work has also been motivated by the hypothesis that physical displacement of a dielectric, as in a Gordon coupler, to match microwave power incident on an EPR sample resonator will result in reduced microphonics compared with use of a metallic slug or screw.

Based on results of Ref. [27], a Rexolite wedge with a reduced-size waveguide dimension *a*_*w*_ = 12.65 mm was chosen. This dimension is calculated by matching the real wave impedance *E*/*H* of the TE_10_ mode between the WR90 waveguide and the Rexolite-filled reduced-size waveguide while keeping the short waveguide dimension *b* fixed. This wave impedance condition was found to minimize the standing waves in the wedge [27]. For this dimension, *γ*^−1^ = 6.73 mm. The Gordon coupler structure was simulated with the DRs and radiation shield. Dimensions are given in Table II. The coupler was found to work with no iris, i.e., the reduced-size waveguide opens directly into the 31 mm shield. Loaded resonator *Q*-values between 1,000 and 60,000 were observed to be critically coupled by varying the position of the wedge from 8 mm to 30 mm. Frequency pulling was only 5 MHz. A square 6 mm iris placed at the boundary between the metallic wall of the shield and the reduced-size waveguide was found to increase the minimum critically coupled resonator loaded *Q*-value to 6,000. With the iris, the electromagnetic mode inside the resonator was more symmetrical and the frequency pulling was smaller. Both the sapphire and rutile DRs were found to be able to be critically coupled over a wide *Q*-value range using the same radiation shield and coupler.

**Table II:** Fabricated shield and Gordon coupler dimensions.

| description | dimension (mm) |
| --- | --- |
| shield diameter and length | 31 |
| iris width and length | 6 |
| iris (wall) thickness | 0.5 |
| waveguide short dimension $b$ | 10.16 |
| WR90 long dimension | 22.86 |
| reduced-size waveguide long dimension $a_w$ | 12.65 |
| reduced-size waveguide length | 30 |
| taper length | 40 |

## V. RESONATOR FABRICATION

Materials for the DRs were sourced, procured, and fabricated by Insaco, Inc. (Quakertown, PA). Synthetic HEM^®^ grown, grade 1 LAOS^®^ sapphire from Crystal Systems, LLC (Salem, MA), was used for the 3–4 μL sample size. This type of sapphire has the highest purity and the lowest number of crystal defects. For the 200 nL sample size, the synthetic SC rutile was purchased from Roditi International Corporation Ltd (London, England) and grown by OXIDE Corporation (Hokuto, Japan). The DRs were ground and then polished to an optical finish including the sample tube hole. The specified dimensions are shown in Table III. For the sapphire, fabricated tolerances were held to within 2%.

For the SC rutile, the ID could not be made smaller than 0.81 mm and, according to finite element simulations, this raises the resonance frequency by 62 MHz. For the SC rutile, the grinding of the OD caused the final shape to become an elliptical cylinder with an OD 2.667 ± 0.032 mm. Finite element simulations showed that this shape causes frequency shifts of between −51 MHz and +1 MHz, depending on the whether the major axis of the ellipse is parallel or perpendicular to the c-axis of the crystal, the angle of which was unknown but is negligible for practical application. The other tolerances were held to within 0.5%.

**Table III:**
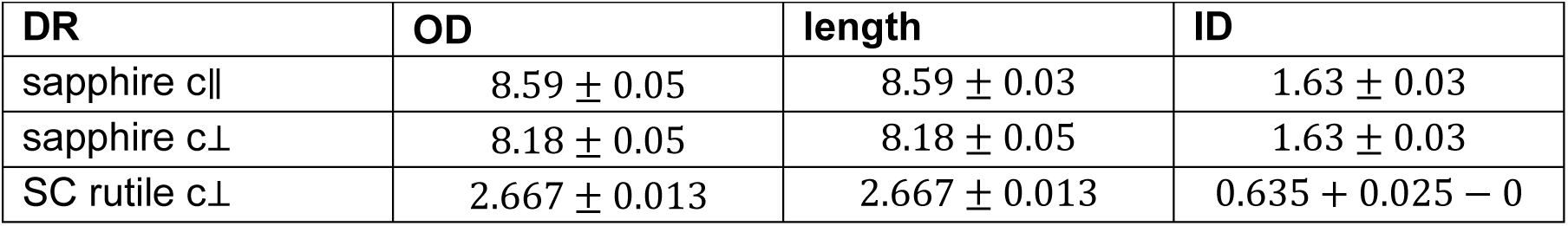
Specified DR dimensions and tolerances (mm)

The sample cells were fabricated by additive manufacturing using the HTL photopolymer, which was selected due to favorable dielectric properties and a lack of dielectric EPR signal, and a Boston Micro Fabrication (Maynard, MA) model S140 3D printer with printing resolution as low as 10 µm, which allowed successful printing of the 50 µm channels in the Aquastar sample cell (Fig. 2B).

**Figure 2:**
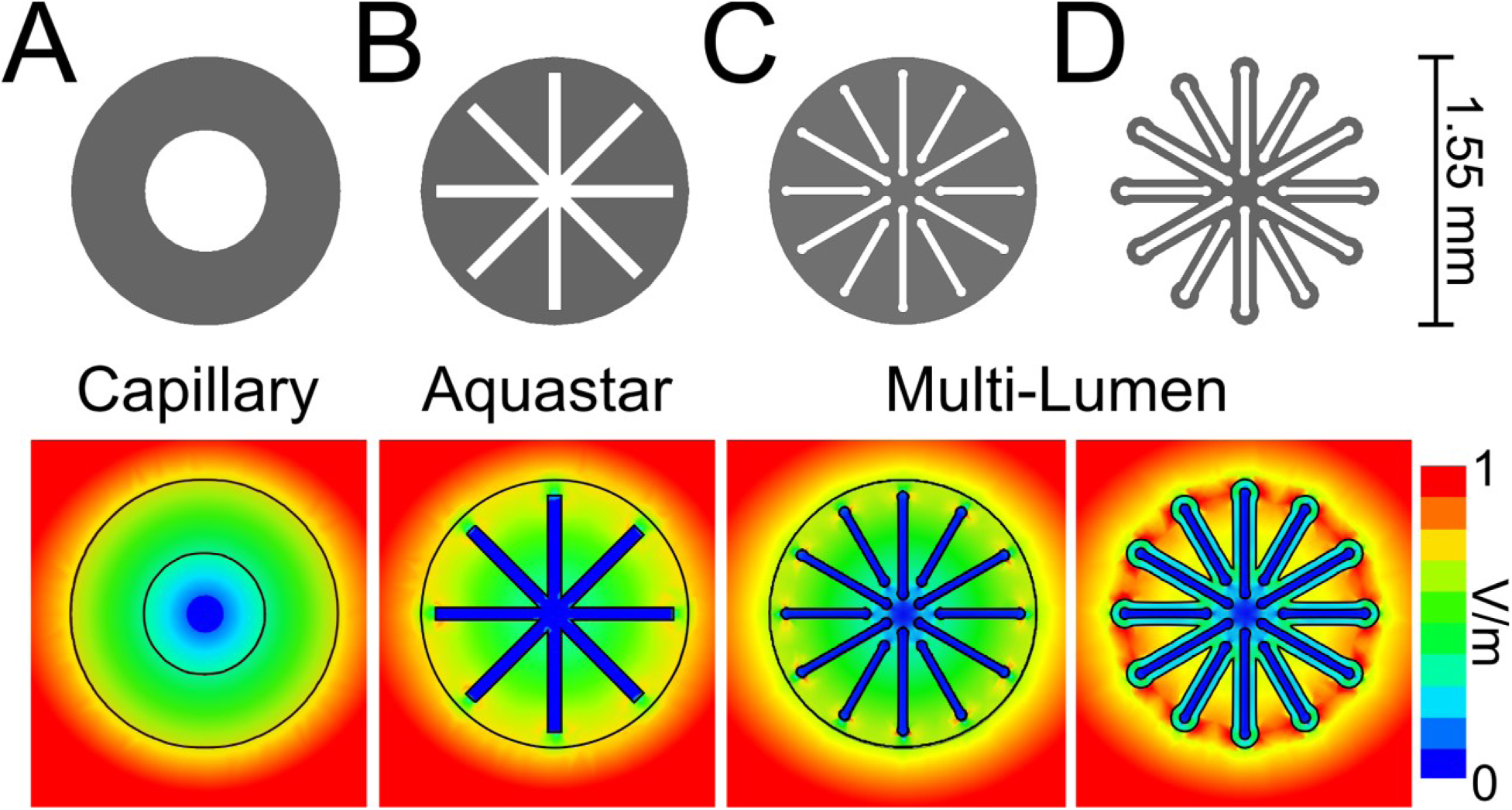
Sample cell geometries (top row) and corresponding normalized electric field magnitude (bottom row) for use with the sapphire resonator discussed in this work. (A) 3 μL capillary, (B) 0.08 × 1.325 mm^2^ Aquastar, (C) solid multi-lumen geometry with 0.05 mm gaps, and (D) multi-lumen geometry with 0.05 mm gaps and 0.05 mm wall thickness.

The Gordon coupler was fabricated from Rexolite, and the cutoff region of a standard waveguide was modified by inserting two machined brass blocks. To hold the coupler component, a slot was cut in the side of the waveguide and pinned to an external sled that was connected to a screw, allowing for coupler adjustment along the waveguide (Fig. 1).

The shield was fabricated out of tellurium copper to the specifications shown in Table II to house the custom sapphire resonator. Three slots were cut along the center of the shield to allow for penetration of the 100 kHz modulation. The modulation coils were wound around a 3D-printed mandrel to approximately 180 µH inductance and are compatible with most commercial spectrometers.

## VI. MULTICHANNEL AQUEOUS SAMPLE CELLS FOR MINIMUM RF LOSSES

Mett and Hyde [23] reported analytical solutions to Maxwell’s equations for multiple aqueous flat cells in a rectangular TE_102_ cavity with an orientation perpendicular to the electric field nodal plane. Results of this study showed that for equivalent amounts of sample, 3–6 times the EPR saturable signal at X-band could be obtained for many thin flat cells than for a single flat cell in the standard orientation. This perhaps counterintuitive result is caused by water having a self-shielding effect where polarization charge developed on the water surface shields the inside from the outside electric field, which lowers RF power dissipation in the water. The high relative dielectric constant of water permits this, but only when the polarization of the electric field is perpendicular to the surface of the water. Tangential electric fields are continuous across the water surface and can produce RF losses larger than the perpendicular orientation, depending on the thickness of the flat cell. A similar effect occurs in metallic conductors where the tangential electric fields cause currents that produce ohmic dissipation, but the perpendicular electric fields do not [26]. A subsequent study by Sidabras et al. [24] added the effects of a PTFE sample cell and confirmed the analytical solutions with finite element simulations, resulting in a U.S. patent [25]. It was found that the sample cell must have a low dielectric constant to produce significant EPR signal enhancement. Later, the team investigated other sample cell shapes for use in LGRs and cylindrical TE_011_ cavities [21].

The self-shielding effect for flat cells in a DR can be determined as follows: Near the DR cylinder axis, the electromagnetic mode has an RF magnetic field directed parallel to the axis and a maximum relatively constant magnitude. The corresponding RF electric field is zero on the cylinder axis, directed azimuthally around the axis, and increases linearly with distance from the axis. For a perpendicular orientation, the flat cells should be placed radially around (or across) the cylinder axis, e.g., as shown in Figs. 2B–D and 3B. The integral form of Faraday’s law can be applied using a circle of radius *r* centered on the cylinder axis. The emf around the loop is equal to the magnetic flux through the loop. Taking the magnetic flux to be the same for the case where there is a single dielectric region and three-layered dielectric regions,

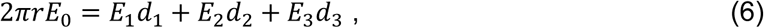

where the *E*s represent the azimuthal electric field and the *d*s the total circumferential arc length of the circle made up by each respective dielectric type, and *d*_1_ + *d*_2_ + *d*_3_ = 2*πr*. The perpendicular electric field boundary condition implies that *∈*_*r*1_*E*_1_ = *∈*_*r*2_*E*_2_ = *∈*_*r*3_*E*_3_, where *∈* is the relative dielectric constant. Taking *∈*_*r*1_ = 1 (air) we find

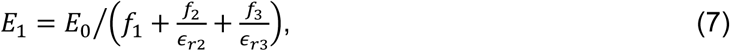

Where 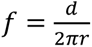 is the fraction of circumference taken up by each dielectric. We also have *E*_2_ = *E*_1_/*∈*_*r*2_ and *E*_3_ = *E*_1_/*∈*_*r*3_. Because the term in braces in Eq. (7) is less than 1, taking region 2 as the sample cell and region 3 as water, 1 < *∈*_*r*2_ ≪ *∈*_*r*3_, Eq. (7) implies that the electric field in air is increased compared with *E*_0_ by the presence of the other dielectrics. But also, the presence of air can reduce *E*_2_ and, because of the large *∈*_*r*3_, will significantly reduce *E*_3_. This analytic result is new and is reflected in the finite element simulations of Figs. 2 and 3. If there is no air, *f*_1_ = 0,

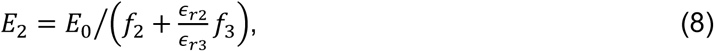

**Figure 3:**
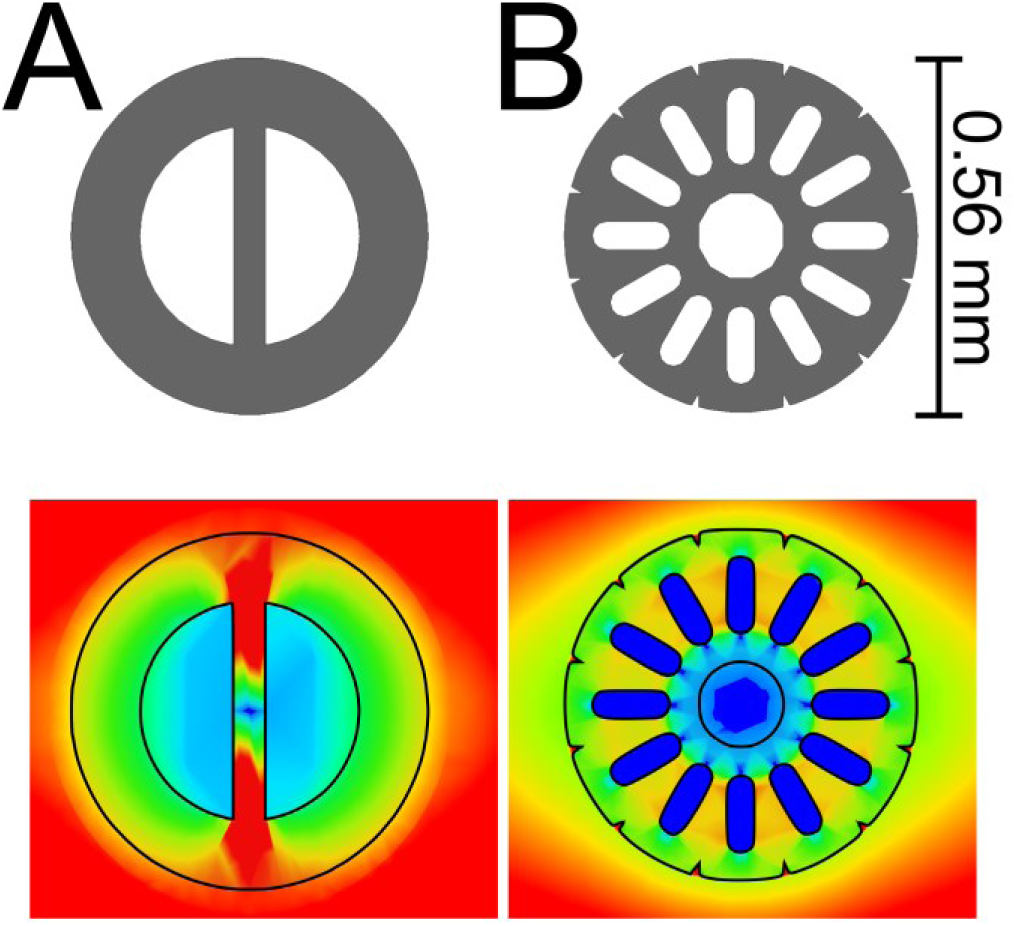
Sample cell geometries (top) and corresponding normalized electric field magnitude (bottom) for use with the rutile resonator discussed in this work. (A) 200 nL single lumen Θ geometry, (B) a multi-lumen geometry.

and *E*_3_ = *E*_2_/*∈*_*r*3_. Again, the term in braces is less than 1 and so the electric field in the dielectric is increased compared with *E*_0_ because of the water. The electric field in the water is reduced, but only by at most a factor of 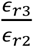 instead of *∈*_*r*3_ when there is also air. This effect is seen in comparing the electric fields in Fig. 2C with 2D. Equation (8) is analogous to Eq. (3) of [24], which is in rectangular geometry. Although these results do not account for the change in the radial wavenumber caused by different dielectrics inside the emf loop, this effect was found to be negligible inside the sample hole of a DR where the radial wavenumber is very small or imaginary in the case of rutile [20].

In Ref. [23], three categories of power dissipation were identified in aqueous flat cells: Type I, caused by RF electric fields tangential to the sample surface; Type II, caused by RF electric fields perpendicular to the surface; and Type III, caused by dipole-like RF electric fields near the ends of the flat cell. Each of these has a different scaling with the flat cell thickness *a*. Because the tangential electric fields tend to circulate around the middle of each individual flat cell, Type I losses scale as *a*^3^. Type I losses unavoidably exist in the perpendicular orientation and typically dominate the total dissipation. Since the perpendicular electric fields are constant across the flat cell, type II losses scale as *a*. The self-shielding of water makes this type of loss negligible unless the flat cell is placed in a region of high RF electric field. Type III losses were also found to be negligible compared with the other two and scale as *a*^4^. By reducing the sample thickness and increasing the number of flat cells for constant sample volume, Type I loss is reduced, and Type II loss remains the same. For a given sample volume, sample losses can be made negligible compared with the losses in the DR itself. It was found that for a given sample thickness, there is an optimum number of flat cells and for a given number of flat cells there is an optimum thickness. This is related to clustering of the flat cells around the electric field node. A similar finding was made in Ref. [26] relating to ohmic losses in conductors with thicknesses comparable to or smaller than an RF skin depth.

We found that the placement of a cylinder of aqueous sample centered on the electric field node with a diameter greater than that of a flat cell thickness always produced higher dissipation than a radial array of flat cells that did not intersect the electric field node. Similarly, an array of intersecting flat cells with water in the center also produced higher dissipation than the non-intersecting radial array, e.g., Fig. 2B compared with 2C and 2D. The presence of an air hole in the center of the sample cell dielectric vs. filled with sample cell dielectric (e.g., Fig. 3B) was found to make a negligible difference.

The constraint of having identically shaped channels is contrary to achieving the lowest sample losses, e.g., compare Fig. 2B with 2C or Fig. 3B. A consideration of fluid flow led us to conclude that placing on each end of the flat cells an enlarged cylinder with a diameter about 33% larger than the flat cell thickness can produce uniform flow rates in the flat cells of arbitrary width. Such bulbed ends were also found to lead to moderately lower RF dissipation in the sample because the bulbs permit a smaller overall sample tube diameter.

A consideration of the Reynolds number for the flow of water in small channels shows that the flow will be laminar if the velocity of the flow is less than about 5 cm/s for the channel thicknesses of 0.05 mm. The flow rate limit proportionally increases with smaller channels. Flow profiles for laminar flow are simple analytical parabolic profiles for flow between two parallel plates and for a circular pipe [36]. For faster flows it is inviscid, and the profile becomes more square. In comparing the laminar flow between two parallel plates and a circular pipe, one can equate the mean fluid velocity between the two for the same pressure drop, and this results in the pipe radius being 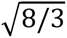 of the plate separation half-distance. One can alternatively equate the maximum velocity in the two for the same pressure drop, and this results in the pipe radius being √2 of the plate separation half-distance. Consequently, it is theoretically possible to have a channel with bulbed ends that produces uniform flow, independent of the channel width. These ratios will be smaller (but always bigger than unity) because the bulbed end on a channel is not a full circle. The exact ratio between the bulb radius and the channel half separation distance can be found by finite element modeling. For the bulbed examples shown in Fig. 2CD, the ratio of 4/3 was chosen, aiming for uniform peak flow velocity.

Simulations of many different multi-channel cells resulted in the designs shown in Figs. 2 and 3. A cell thickness of 0.050 mm was deemed able to be readily fabricated. Table III shows a comparison of the performance of these cells with each other and with a capillary in the same DR hole size as Table I. PEC was used for the shield to highlight the losses in the sample, sample cell, and dielectric. The ratio of the sample power losses *P*_*s*_ to the total power loss *P*_*l*_, and the ratio of the sample cell power loss *P*_*h*_ to the total is also shown. One can see the *Q*, *Λ*, and *S*_*s*_ are all significantly higher for the multi-lumen samples than the capillary. This is because the sample losses are reduced. The losses are reduced further with a greater number of smaller channels. It is possible to reduce the sample losses to a small fraction of the DR losses by making the thickness of the sample flat cell sufficiently small while increasing the number of cells to keep the sample volume constant. Introducing layers of air between channels at larger radii significantly reduces both the sample and sample cell losses, even for the HTL. Notice that the highest *Q* and *Λ* in Table III are slightly smaller (due to sample) than those shown in Table I (for PEC), and ratios of these can be used to estimate *S*_*s*_ for other configurations. To ensure simulations were accurate and to account for the losses associated with the Rexolite resonator holders shown in Fig. 1, the loss tangent of the sapphire was slightly increased to better match the measured *Q*-value. This adjusts the simulated Λ-values to more practical values.

Color plots of the RF electric field magnitude in the vicinity of the aqueous sample and sample cell from the finite element simulations for configurations reported in Table II are shown in Fig. 2A–D for sapphire and Fig. 3AB for rutile. The capillary plot in Fig. 2A shows continuity of the tangential electric field across the sample surface and the cause of the relatively large sample losses. The azimuthally solid sample cells of Figs. 2B,C and 3B can be compared with the same sample configuration with the introduction of azimuthal air gaps between channels of Fig. 3B or 3C, which show a significant reduction in electric field in the sample and sample cell, resulting in a higher EPR signal intensity for sapphire and rutile, as outlined in Table III and Table IV, respectively. Within Table III is also the simulated EPR saturable signal for a 2.2 μL two-loop– one-gap LGR that was previously commercially available from Molecular Specialties, which had become a standard for power saturation CW experiments due to its relatively large Λ-value and concentration sensitivity.

**Table IIIa:**
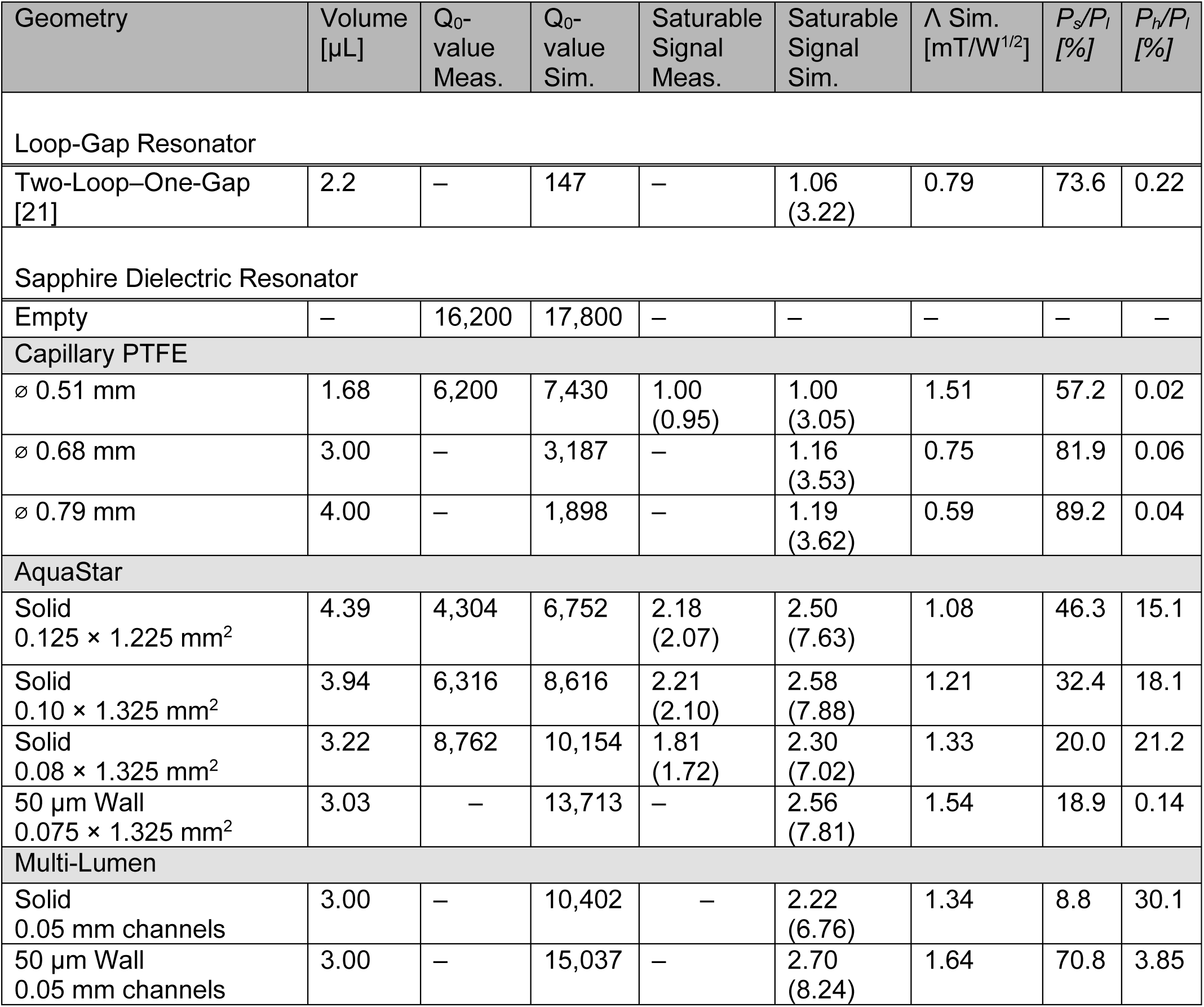
Simulated and measured EPR signal intensities for several sample geometries.

**Table IV:** Simulated characteristics of several sample tube types for the rutile resonator.

| Geometry | Volume<br>[$\mu\text{L}$] | Q<sub>0</sub>-<br>value<br>Sim. | Saturable<br>Signal<br>Sim. | $\Lambda$ Sim.<br>[mT/W<sup>1/2</sup>] | $P_s/P_l$<br>[%] | $P_h/P_l$<br>[%] |
| --- | --- | --- | --- | --- | --- | --- |
| Rutile Dielectric Resonator (c-axis $\perp$) | | | | | | |
| Empty | – | 17,800 | – | – | – | – |
| Capillary PTFE |  |  |  |  |  |  |
| ∅ 0.31 mm | 0.200 | 5,011 | 1.00<br>(2.19) | 6.87 | 94.1 | 0.05 |
| Multi-Lumen |  |  |  |  |  |  |
| Θ geometry | 0.200 | 7,482 | 1.45<br>(3.17) | 9.77 | 79.1 | 9.6 |
| 12-star bulb<br>0.05 mm channels | 0.200 | 8,929 | 2.35<br>(5.14) | 13.95 | 53.5 | 25.2 |

## VII. EXPERIMENTAL EPR CHARACTERIZATION

Listed in Table III are bench and EPR measured characteristics of the sapphire DR with Aquastar sample cell geometries (Fig. 2B) with fin widths of 125, 100, and 80 μm corresponding to 4.4, 3.9 and 3.2 μL volumes, respectively. Current internal feature limitations on our in-house 3D printer made the smaller fin widths and geometries shown in Figs. 2CD and 3 challenging. Sample cells with a standard cylindrical cavity and the Aquastar geometry were bench characterized and tested experimentally for comparison to the simulated data. Power saturation CW EPR spectra were recorded at X-band on an ELEXSYS 500 spectrometer (Bruker; Billerica, Massachusetts) at room temperature operating with the following parameters: 345.4 mT center field, 10 mT scan width, variable incident microwave power, 5.12 ms conversion time, 1.28 ms time constant, 0.1 mT modulation amplitude, 100 kHz modulation frequency, 41.92 second sweep time, and 9.69 GHz microwave frequency.

As shown in simulations and summarized in Table III, there is little improvement by increasing the volume of a PTFE capillary in a sapphire resonator due to the significant reduction in the *Q*-value as volume increases. Therefore, practically, the best performance of a capillary sample tube in our spectrometer was found when the *Q*-value of the sapphire resonator with a 1.67 μL active volume was used. The power saturation CW EPR data were plotted using Graphpad Prism, resulting in Fig. 4. The PTFE cylindrical sample cell has an active volume of 1.67 µL and the power saturation data has been normalized to the peak EPR signal, which is referred to as the saturable signal. Saturable signal is defined as the EPR signal obtained at constant RF magnetic field incident on the sample.

**Figure 4:**
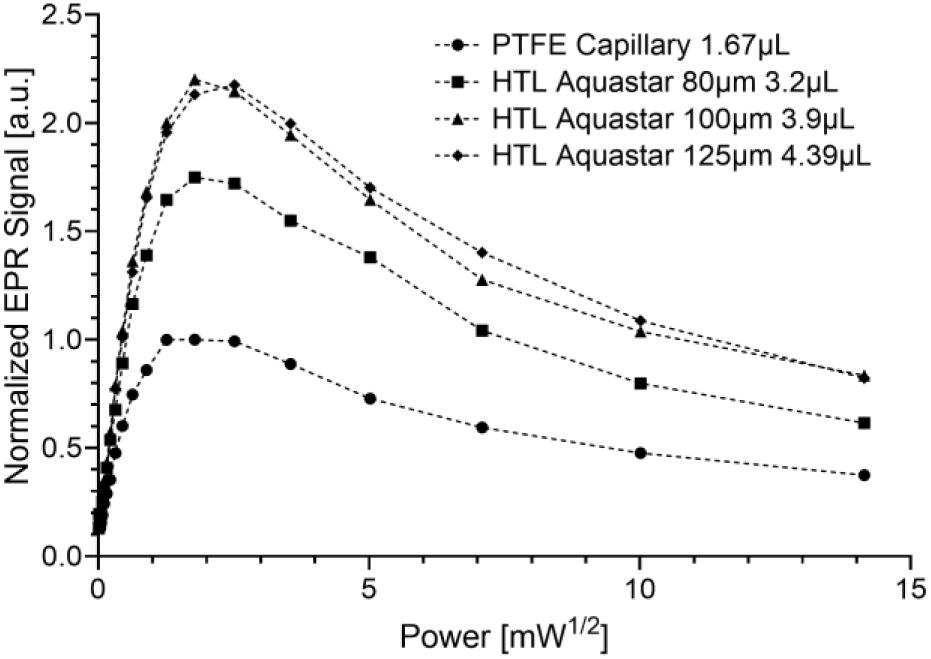
Power saturation curves for the data tabulated in Table III. Data are normalized to the 1.67 μL PTFE capillary maximum signal (saturable signal). Three 3D printed Aquastar geometries were tested with varying fin width and total volume.

Using the Boston Micro Fabrication 3D printer, HTL sample tubes with the Aquastar geometry and with a 125 μm fin width were fabricated (active volume of 4.35 µL). This Aquastar geometry was recently used experimentally in Ref. [37]. In comparison with the capillary, a 2.18-fold signal improvement was measured for this multi-lumen geometry. To decrease the required sample volume, we then 3D printed an HTL Aquastar geometry with a 100 μm fin width (active volume of 3.9 μL). Despite the 20% reduction in volume, the EPR saturable signal intensity remained nearly the same as the 4.35 µL Aquastar, with a signal improvement of 2.21-fold compared with the capillary. A further reduction in the volume by reducing the fin width to 80 μm only produced a 1.81-fold increase in signal compared with the capillary; this is in contrast with the finite element modeling simulations that predicted a 2.3-fold increase. This discrepancy with the 80 μm fin width is due to challenges printing internal features. Although the Boston Micro Fabrication 3D printer can easily print 10 μm features on a surface, printing features within a volume is far more challenging. Several key optimizations are required before further reduction of fin width is feasible: *(i)* Temperature of the HTL material while printing. Due to the viscosity of the material, varying the temperature from the current room temperature printing to 40 °C may help with internal features due to resin staying within the feature causing reduction of actual fin width. *(ii)* Printing power and time. Currently we are using an incident power and curing time per layer that maximizes printing success, but this may be too long, resulting in overcuring of the cross-sectional geometry of the sample cells. Any overcuring would result in reduced accuracy of internal features. Our current protocol is optimized for producing 16 sample cells within 8 hours. *(iii)* Cleaning and post-printing curing. We have created a rigorous cleaning procedure using isopropyl alcohol flow and sonication. After cleaning, the sample cells are placed in a Formlabs (Formlabs Inc., Somerville, MA) Form Cure, which exposes the sample cells to a 405 nm light at room temperature. This final curing stage was shown to significantly increase (30+%) the *Q*-value of the resonator when compared with uncured sample cells. Due to the large aspect ratio of the sample cells (1.55 mm wide by 35 mm long) the curing must be performed in a 2 mm quartz capillary to reduce warping of the sample cells. When cured in the 2 mm quartz capillary, no bending of the sample cells is evident.

In simulation, a study to further reduce the sample volume to 3 μL was performed by assuming we could print 75 μm fins along with reducing the HTL plastic around the sample while maintaining a 50 μm wall. By reducing the lossy HTL plastic from the sample cell geometry, a similar simulated increase (approx. 2.5-fold) to the 4 μL sample would be available despite the 25% reduction in volume.

Further simulation studies using the multi-lumen geometry of Fig. 2CD show that despite the significant reduction in electric field within the sample by using 50 μm sample channels the losses associated with the HTL plastic dominate, resulting in no notable improvement compared with the Aquastar geometry. However, reducing the HTL plastic down to the 50 μm wall would result in a 2.7-fold increase of EPR signal intensity compared with the capillary.

Finally, presented in Table IV, are the simulations of the SC rutile DR with the HTL geometries shown in Fig. 3 compared with a PTFE capillary with a volume of 200 nL. As shown in Fig. 3A and Table IV, the Θ-geometry increases the EPR signal intensity by a factor of 1.45 while maintaining a very simple printable geometry. This is due to the creation of a perpendicular surface within a capillary, reducing the electric field and breaking up the tangential electric field component, Fig. 3C. However, with the 12-star bulb geometry shown in Fig. 3B, a simulated increase by a factor of 2.35 is possible, requiring the printing of 50 μm channels.

It should be noted that the simulated values, given in parentheses, of the saturable signal can be directly compared between Table III and Table IV. In doing so, it is found that the 1.67 μL capillary within the sapphire will have similar EPR signal intensity compared with the Θ-geometry within the rutile, 3.05 and 3.17, respectively. The significant increase in RF magnetic field due to the dielectric constant incident on the sample, along with the high *Q*-value maintained by breaking up the tangential electric field, makes up for the 8.4-fold decrease in sample volume.

## VIII. CONCLUSIONS

We present here new innovations in resonator and sample cell designs that reduce the volume required per sample while improving spin sensitivity. Advances in 3D printing technology have enabled the printing of creative sample cell geometries that were previously unattainable due to constraints with extrusion techniques that required continuous openings and larger dimensions that limited these advances. With printing technologies of 10 µm resolution, we were able to print in-house the Aquastar design with 3–4 µL volumes. Printers with resolution of 2 µm are now commercially available and, with further optimization, will enable the future printing of the designed 200 nL sample cells and more complex 3 μL samples. Resin selection is also an important factor as RF losses and EPR signals need to be minimized to be effective. Many resins exist, but most have an intrinsic EPR signal (e.g., Rodgers radix). Fortunately, HTL does not exhibit an EPR signal and enabled the experimental studies presented here. Future advances in resins will also enable future developments in sample cell designs.

In summary, we have shown that complex sample cell geometries beyond a cylindrical cavity allow novel low volume spin sensitivity. In addition, the use of sapphire and rutile DR designs with a Gordon coupler further enables signal enhancement for low volume biomedical EPR samples.

## IX. ACKNOWLEDGMENTS

The authors thank Insaco, Inc. (Quakertown, PA), for the fabrication of the DRs, particularly the SC rutile, which required the development of new methods, and Boston Micro Fabrication (Maynard, MA) for 3D printer support. This work was supported by the National Institutes of Health grants GM140385 (to CSK and MTL) and GM149568 (to JWS).

## AUTHOR DECLARATIONS

## Author Contributions

**<u>R. M. Mett:</u>** Conceptualization (lead); Investigation (equal); Methodology (equal); Visualization (equal); Writing – original draft (lead)

**<u>A. M. Garces:</u>** Investigation (supporting)

**<u>A. Anilkumar:</u>** Investigation (supporting)

**<u>J. T. Wehrley</u>:** Investigation (supporting)

**<u>M. T. Lerch:</u>** Funding acquisition (equal); Project Administration (equal); Writing – review and editing (supporting)

**<u>B. S. Klug:</u>** Funding acquisition (equal); Methodology (supporting); Project Administration (equal); Writing – review and editing (lead)

**<u>J. W. Sidabras:</u>** Conceptualization (equal); Investigation (equal); Methodology (equal); Project Administration (equal); Supervision (lead); Visualization (lead); Writing – review and editing (lead)

## DATA AVAILABILITY

The data that support the findings of this study are available from the corresponding author upon reasonable request.

## Footnotes

1 If the RF magnetic field is linearly polarized, 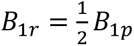.

2 An equation similar to Eq. (2) was reported by Tschaggelar et al. [18]; however, a homogeneous *B*_1_ field over the resonator was assumed. From Eq. (2), it follows that the *B*_1_ value in Eq. (10) of [18] is quantitatively a root mean square value when *Q* is replaced by the critically matched loaded *Q*-value *Q*_*L*_ = *Q*/2.

